# SpaNorm: spatially-aware normalisation for spatial transcriptomics data

**DOI:** 10.1101/2024.05.31.596908

**Authors:** Agus Salim, Dharmesh D Bhuva, Carissa Chen, Pengyi Yang, Melissa J Davis, Jean YH Yang

**Affiliations:** Melbourne School of Population and Global Health, The University of Melbourne, Melbourne, VIC, 3010, Australia; Bioinformatics Division, Walter and Eliza Hall Institute of Medical Research, Parkville, VIC, 3052, Australia; School of Mathematics and Statistics, The University of Melbourne, Melbourne, VIC, 3010, Australia; Baker Heart and Diabetes Institute, Melbourne, VIC, 3004, Australia; South Australian ImmunoGENomics Cancer Institute, Faculty of Health and Medical Sciences, The University of Adelaide, Adelaide, SA, 5005, Australia; Precision Cancer Medicine, South Australian Health and Medical Research Institute (SAHMRI), Adelaide, SA, 5000, Australia; School of Medical Sciences, Faculty of Medicine and Health, The University of Sydney, Sydney, NSW, 2006, Australia; Computational Systems Biology Unit, Children’s Medical Research Institute, Westmead, NSW, 2145, Australia; Sydney Precision Data Science Centre, The University of Sydney, Sydney, NSW, 2006, Australia; School of Biomedicine, Faculty of Health and Medical Sciences, The University of Adelaide, Adelaide, SA, 5005, Australia; Isomorphic Labs, London, United Kingdom; School of Mathematics and Statistics, The University of Sydney, Sydney, NSW, 2006, Australia; Charles Perkins Centre, The University of Sydney, Sydney, NSW, 2006, Australia

## Abstract

Library size normalisation is necessary to enable comparisons between observations in transcriptomic datasets. Numerous methods have been developed to normalise these effects with sample and gene specific adjustments. However, in spatial transcriptomics data, normalisation is complicated by the fact that spatial region-specific library size confounds biology. The most popular approach of adapting methods developed for single-cell RNA-seq data has been shown to excessively remove biological signals associated with spatial domains and thus results in poorer downstream domain identification. To this end, we propose the first spatially-aware normalisation method, SpaNorm. SpaNorm concurrently models spatial library size effects and the underlying smooth biology, to tease apart these effects, and thereby remove library size effects without removing biology. This is achieved through optimal decomposition of spatially smooth variation into those related and unrelated to library size and the use of location-specific scaling factors. Using 27 tissue samples from 6 datasets spanning 4 spatial platforms, we show that SpaNorm outperforms current state of the art methods at retaining biological information in the form of spatial domains and spatially variable genes (SVGs) better than 4 commonly used single-cell normalisation approaches. SpaNorm is versatile and it can be used for both spot-based and subcellular spatial transcriptomics data. Notably, the benefit of using SpaNorm is more pronounced for the latter data such as those from Xenium, STOmics and CosMx platforms for which the proportion of genes exhibiting region-specific library size effect is higher. SpaNorm works equally well with segmented cell-level data and spot-based data where each spot contains multiple cells.

## Introduction

Advances in spatial profiling technology have transformed our comprehension of multicellular biological systems. The emergence of both spot-based spatial transcriptomics technologies (ST) such as 10x Genomics Visium [1] as well as subcellular spatial transcriptomics (SST) technologies, such as 10x Genomics Xenium [2], NanoString CosMx [3], BGI Stereo-seq [4], and Vizgen MERSCOPE [5], hold the promise to address previously inaccessible biological questions and enhance our understanding of intercellular communication by preserving tissue architecture. While these innovative spatial transcriptomics technologies offer the potential to uncover new insights into regional variations in cell density and composition, the challenge of effectively removing varying library sizes across regions (Fig. 1A and 1B) hinders our ability to detect spatial variation signals from the data. This can potentially impact downstream analyses such as clustering, regional segmentation and identification of spatially variable genes (SVGs).

**Fig. 1.**
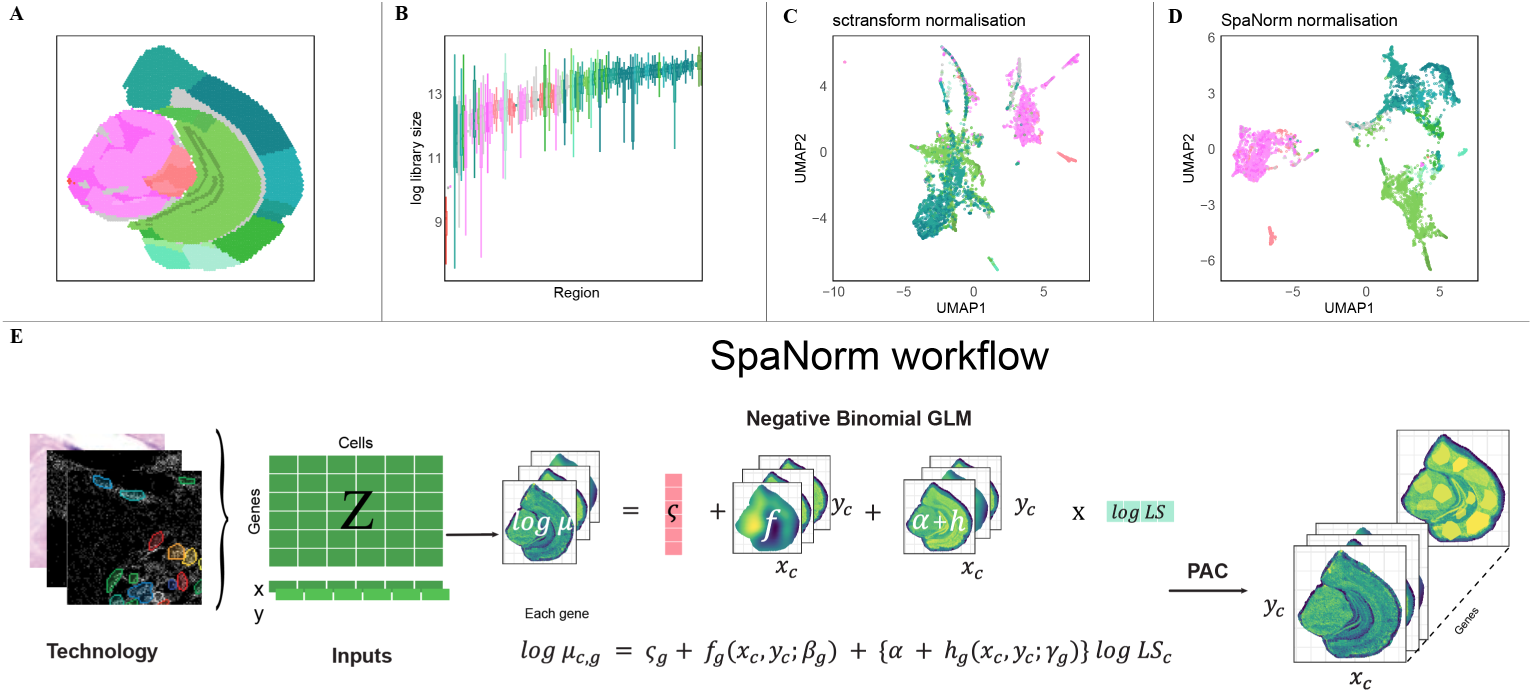
(A) Spatial regions in STOmics Human Brain Dataset (B) Library size distribution differs across regions, with WM (white matters) region has lower library size compared to other regions. (C) UMAP of sctransform adjusted data showing clear separation of spatial regions. (D) UMAP of SpaNorm adjusted data showing less distinct separation of spatial regions. (E) SpaNorm workflow: SpaNorm takes the gene expression data and spatial coordinates as inputs. Using genewise Negative Binomial (NB) model, SpaNorm decomposes spatially-smooth variations into those unrelated to library size (LS), representing the underlying true biology and those related to libray size. The adjusted data is then produced by keeping only the variation unrelated to library size.

Currently, the removal of library size effects from spatial transcriptomics data is under debate. Those in favour would argue that total molecule counts represent technical unwanted variation from imperfect molecules capture and would typically use methods originally developed for single-cell RNA-seq (scRNA-seq) data [6–8] that ignore the spatial information. Many of these library size normalisation approaches use global scaling factors and may suffer when the data are confounded by spatial region-specific library size biases. In particular, these normalisation methods tend to remove signals associated with the spatial domain (Fig. 1C), and have led to arguments that library size normalisation should not be performed prior to downstream analyses [9] or at least prior to spatial domain identification unless addressed using methods that take spatial information into account [10]. To this end, there is a need for normalisation techniques that leverage spatial information to eliminate this region-specific library size bias while retaining biological signals for downstream analyses as effective library size normalisation can improve spatial domain identification and other downstream analyses.

Here, we develop SpaNorm, a normalisation method that utilises spatial information and gene expression simultaneously, allowing optimal identification of spatial domains (Fig. 1D) and spatially variable genes (SVGs). We achieve this through three key innovations: (1) optimally decomposing spatially-smooth variation into library size associated and library size independent variation via generalized linear model (GLM); (2) computing spatially smooth location- and gene-specific scaling factors; and (3) using percentile-invariant adjusted counts (PAC) [11] as normalized data for downstream analyses. Fig. 1E) provides a detailed overview of the SpaNorm approach.

## Results

### Library size effects are region-specific in spatial transcriptomics data

We first establish evidence that library size effects vary across spatial domains. Comparing models with global and region-specific library size effects, we estimate the proportion of genes that exhibit spatial variations in their library size effects (see Material and Methods section). From Fig. 2A, we can infer that the proportion of genes with region-specific library size effects varies from around 25% to almost 100% across datasets. Overall, Xenium and STOmics datasets have the highest proportion of genes with region-specific effects, followed by CosMx dataset, and finally the Visium dataset. To demonstrate that these results are not due to our manual region annotations, we performed a sensitivity analysis where each dataset was split into rectangular grids and estimated the proportion of genes that exhibit grid-specific library size effects. For the majority of the datasets, the results show an even higher proportion of genes that exhibit variation in their library size effects under the grid-based method (Supplementary Figure 1).

**Fig. 2.**
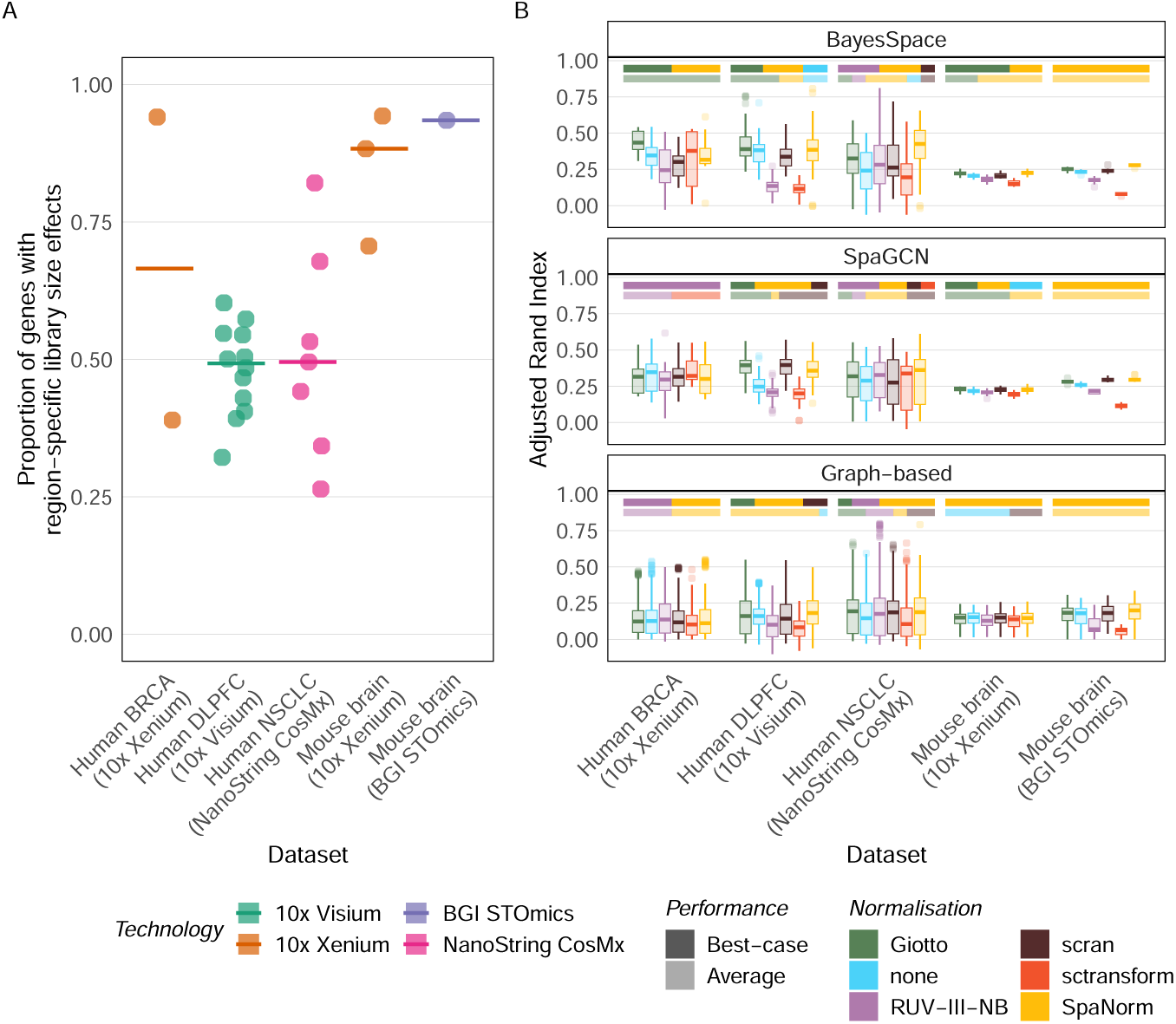
(A) Estimated proportion of genes with region-specific library size effects. On average, CosMx and STOmics datasets have the highest proportion of genes exhibiting region-specific effects, followed by Xenium. Visium datasets have the lowest proportion. (B) Adjusted Rand Index of clusters identified using differently normalised data vs annotated spatial regions. Boxplots show the summary by platform. The coloured bars above each group of boxplots indicate the best-performing method for each dataset that makes up the group, based on maximum (darker-shade) and median (lighter-shade) statistics.

### SpaNorm preserves spatial domain signals

Next, we examine how the spatially-dependent library size effects (scaling factor) in SpaNorm can improve downstream analyses. For this purpose, we first compared SpaNorm to other normalisation methods in terms of their ability to retain spatial domain information. We use the ratio of between-region to within-region variations to measure the strength of signals associated with the spatial domain in each gene. Comparing these ratios for the raw and differently-normalised data (Supplementary Figure 2), we found that SpaNorm retains the most signal (higher ratio), followed by scran and RUV-III-NB while sctransform and Giotto retain the least. Giotto particularly retains less signal for the Xenium datasets (Mouse Brain, ILC, and IDC).

Clustering has been used as one of the main tools to identify distinct spatial regions. To examine how better retention of spatial domain signals translates into improved identification of spatial regions, we benchmarked SpaNorm against several alternative normalisation methods using our previously established benchmark [10]. Three clustering methods: graph-based, SpaCGN and BayesSpace, were applied using a range of parameter settings (see Materials and Methods). Fig. 2B shows that single-cell RNA-seq inspired graph-based methods that use expression data alone have lower clustering accuracy across all platforms compared to the spatially-aware methods BayesSpace and SpaCGN. Furthermore, with graph-based clustering, the choice of normalisation method has little impact on clustering accuracy, with the exception of sctransform where lower performance is observed in the Mouse brain STOmics, Human DLPFC Visium and Human NSCLC CosMx data.

### SpaNorm enables more stable performance

The choice of normalisation methods has a pronounced impact on clustering accuracy with BayesSpace and SpaCGN. When clustering with BayesSpace, which has a slightly better overall accuracy than SpaCGN, we observe that SpaNorm has the best performance (measured using maximum ARI) among Mouse brain STOmics samples and approximately half of Human NSCLC CosMx, Human DLPFC Visium and Human BRCA Xenium samples. SpaNorm also has comparable performance to Giotto for the Mouse brain Xenium when clustering using BayesSpace (Fig. 2B). The second best method is Giotto with maximum ARI in two out of three Mouse brain Xenium samples and one of the BRCA Xenium samples (Fig. 2B). On the other hand, no normalisation (none) rarely produced the best performance across most datasets, highlighting the need for appropriate library size normalisation for downstream analyses of spatial transcriptomics data.

When clustering with SpaCGN, SpaNorm is still the best performing method for the Human DLPFC Visium datasets, and the Mouse brain STOmics datasets. It is joint-best (with Giotto) for the Mouse brain Xenium datasets and is second to RUV-III-NB for the Human NSCLC CosMx dataset. Overall, SpaNorm achieves the most stable relative performance across all datasets across various tissues and platform types (Supplementary Figure 3).

### SpaNorm improves SVG detection and concordance

Beyond spatial domain identification, we show the benefits of SpaNorm normalisation in consistently detecting spatially variable genes (SVGs). We demonstrate performance using simulated datasets where the true SVGs are known, and using sequential replicate sample from real world datasets where SVGs identified should be consistent. For the former experiment, we generated realistic simulated datasets using scDesign3 [12] where 100 genes were designated as true SVGs. Supplementary Figure 4 shows that among the top 100 SVGs identified, SpaNorm consistently calls the highest proportion of true SVGs correctly, followed by sctransform and Giotto, with RUV-III-NB having the lowest concordance.

SpaNorm is also better at detecting true SVGs in real datasets. Fig. 3B and Supplementary Figures 5-6 show the expression of six true SVGs from Xenium Mouse Brain datasets [13]. While Giotto and no Normalisation produce stronger signals for general neuronal subtype markers that distinguish granule neurons in the dentate gyrus (*Prox1*) from pyramidal neurons in CA1-3 (*Neurod6*) [14], SpaNorm produces stronger signals for detecting specific markers of pyramidalneurons in different CA regions, namely *Wfs1* in CA1 region, *Necab2* in CA2 region and *Slit2* in CA3 region (Fig. 3C).

**Fig. 3.**
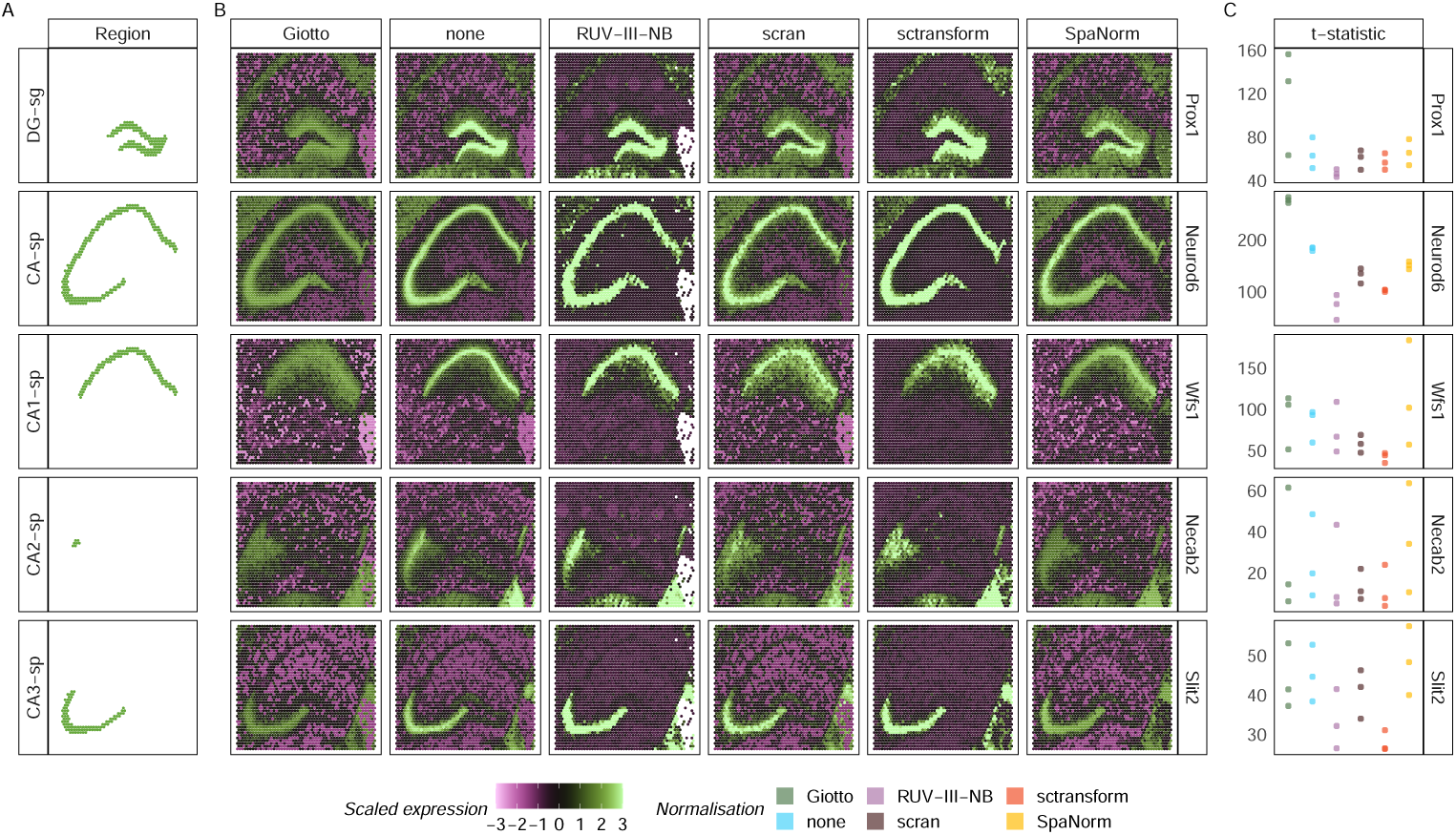
(A) Sub-structures of the mouse hippocampus. (B) Expression of 5 spatially variable genes (SVGs) in Xenium Mouse Brain dataset (replicate 1) under differently normalised data. *Prox1* differentiates the dentate gyrus (DG-sg) from pyramidal layers in CA1-3 (*Neurod6*). *Wfs1*, *Necab2 Slit2* are enriched in CA1-sp, CA2-sp and CA3-sp regions, respectively. (C) t-statistic from three replicates for comparing expression inside vs outside the regions. Higher statistic means stronger spatial signals.

SpaNorm also generally improves the concordance of SVG list. Using replicate datasets (three Visiums, two Xeniums and one CosMx), we compare the concordance of SVG lists among datasets belonging to the same replicate. Fig. 4 shows that no Normalisation results in the highest concordance of SVG list among replicates of Visium datasets, followed by Giotto and SpaNorm. While it also has very high concordance among replicate of the other sets. Further investigation indicates that the top SVGs under no Normalisation is dominated by genes that are stably expressed across cell types (’housekeeping’ genes). The SVG test statistic that measures the strength of signals shows that the signals for these stably-expressed genes are much stronger in the un-normalised data (see Supplementary Figure 7), suggesting that the top SVGs from the un-normalised data reflect library size effects rather than true biological signals. Excluding the un-normalised data, we observe that SpaNorm provides the best concordance for replicates of the two Xenium datasets, while for CosMx replicates, transform and scran are slightly better than SpaNorm. RUV-III-NB consistently has the lowest concordance across all datasets (Fig. 4B).

**Fig. 4.**
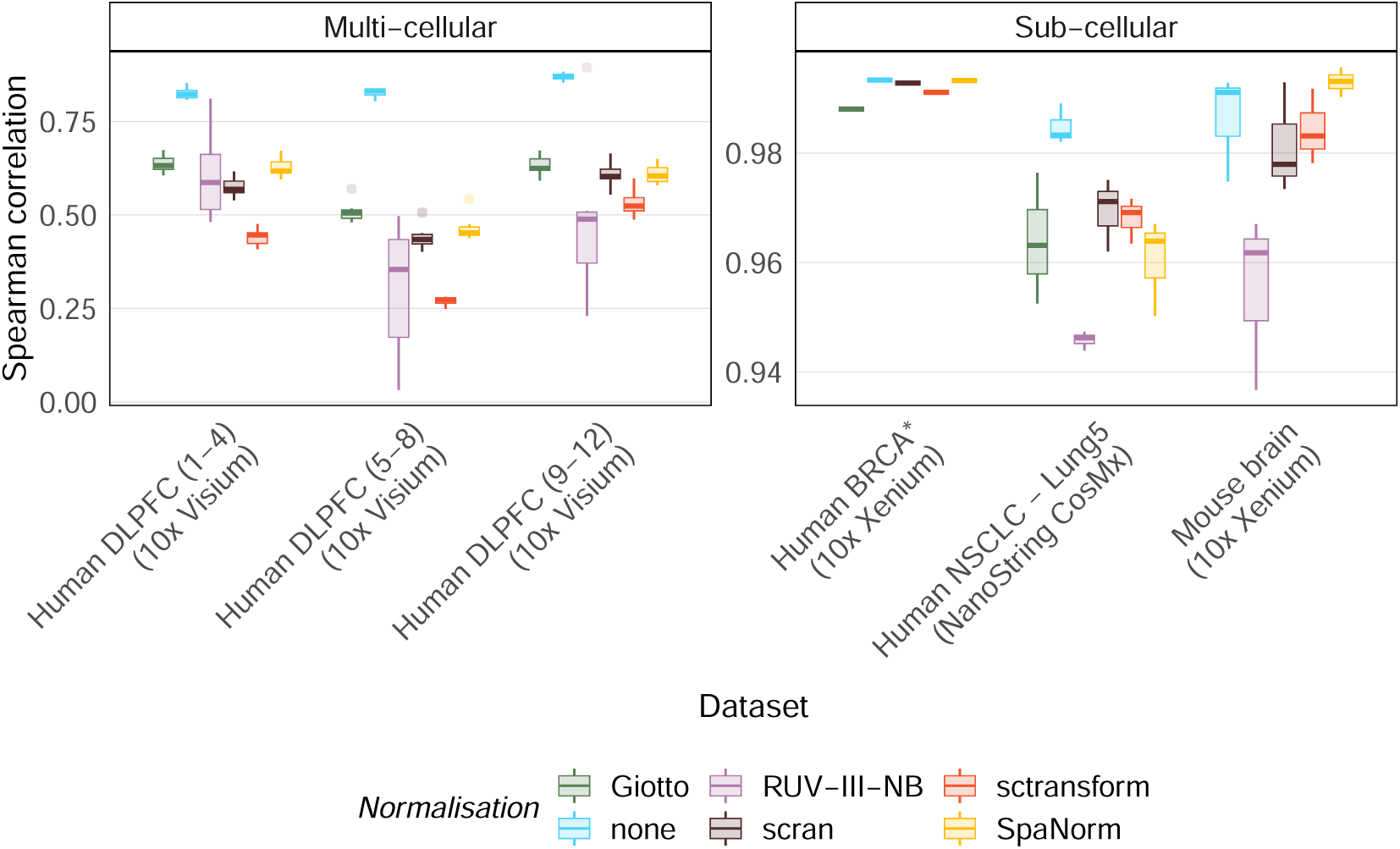
Within-set Spearman’s correlations of MERINGUE’s SVG statistic. (left) Visium Human DLPFC sets, (right) CosMx Human Lung, Xenium Human Breast Cancer and Xenium Mouse Brain sets. Higher Spearman’s correlation indicates more consistent SVG rankings among replicates of the same set. (* Breast cancer dataset with replicates of the same section)

### SpaNorm enhances biological signals from lowly expressed genes

Lower library sizes due to technical effects can make it difficult to detect marker genes that are essential for identifying spatial domains. Though library size normalisation can adjust these effects, lowly expressed genes are still difficult to detect. MOBP is one such marker gene that marks oligodendrocytes that are enriched in the white matter of the human brain (Fig. 5A) [15]. When analysing the 10x Visium human DLPFC datasets, we saw that MOBP was lowly expressed in two of the twelve samples. In these datasets, we saw that the library size of spots from the white matter (WM) had particularly low library sizes (Fig. 5B). Not normalising library size effects would lead to the conflicting conclusion that MOBP was excluded from the white matter (Fig.5C). Giotto, scran, and sctransform were able to detect signals at the boundary of the white matter but not within. Only SpaNorm was able to detect signals both within and at the boundary of the white matter region, however, MOBP detection was relatively weaker at the core of the region. As SpaNorm models the expression of each gene spatially, it enables borrowing of information from surrounding regions, and this can be used to obtain a better region-specific estimate of each gene’s expression. Inspecting the mean estimate of MOBP, we saw that it was significantly higher in the white matter compared to other regions of the tissue (Mean Bio in Fig.5C). This observation was also consistent across other samples from this dataset (Supplementary Figure 8).

**Fig. 5.**
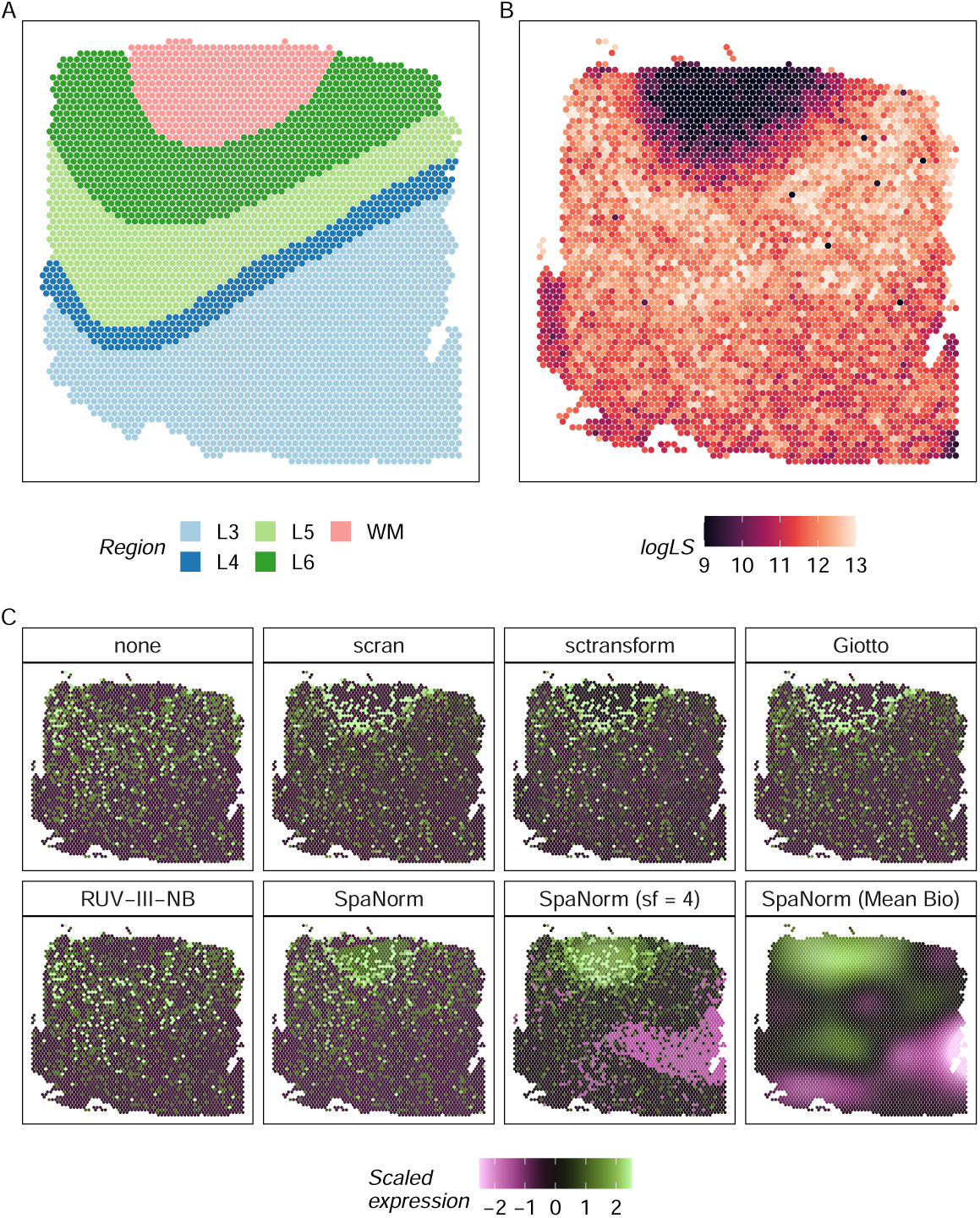
(A) Expert annotated regions in the Human DLPFC dataset. (B) log library size for each spot shows a significant drop in library sizes in the white matter region (WM). (C) Normalisation using different approaches is unable to recover the expression of a marker gene, MOBP, of oligodendrocytes that are enriched in the white matter region. Not normalising results in the contradictory inference of exlucsion of this gene from the white matter. SpaNorm detects this signal as indicated by the mean estimate of this gene, and the effect is more evident when the percentile-invariant adjusted count (PAC) are scaled up by a factor of 4 (sf = 4) during adjustment.

## Discussion and Conclusion

Here we present SpaNorm, a normalisation method that recognises the region-specific nature of library size effects and distributions. Using 27 realworld datasets, we benchmarked SpaNorm against other normalisation methods and demonstrated that SpaNorm is better at retaining spatial domain signals for clustering and detecting true SVGs. SpaNorm’s running time increases only linearly as a function of the number of cells. For the datasets we used in our benchmarking study, the longest running time was around 9 minutes for Xenium Breast Cancer datasets with around 60,000 cells (Supplementary Figure 9).

To maximise the potential of SpaNorm’s normalised data, we recommend using spatially-aware clustering algorithms such as BayesSpace and SpaCGN, for which the comparative advantage of SpaNorm is more pronounced. While SpaNorm can be used for both spot-based and subcellular spatial transcriptomics (SST) data, we observed that the benefit of using SpaNorm is more for SST data such as those from Xenium, STOmics and CosMx platforms for which the proportion of genes exhibiting region-specific library size effect is higher.

For data generated using SST technologies, in order to extract cell-level data, segmentation to detect cell boundaries can be carried out prior to downstream analysis. An alternative is to use grid-based methods [10] whereby no segmentation is performed and instead molecule counts that fall into each grid are simply summed up. Our benchmarking consists of 25 grid-based and 2 segmentation-based datasets. Our empirical evidence shows that SpaNorm’s performance is not sensitive to this decision and the algorithms work equally well for segmentation-based data or grid-based data that consist of counts from multiple cells.

Optimal normalisation of spatial transcriptomics (ST) data has been difficult to achieve because library size effects and distribution are potentially region-specific. These two unique features of ST data do not exist in single-cell RNA-seq (scRNA-seq) data. It is thus not surprising that direct applications of normalisation methods developed for scRNA-seq data often results in the removal of spatial domain signals in addition to removing the library size effects.

SpaNorm currently only deals with library size effect but can be extended to handle other unwanted variation such as ‘batch’ effect introduced when data are acquired through multiple fields of views [16]. Fields of View (FOV) effect introduces discontinuity in the spatial patterns. Since SpaNorm relies on decomposing spatially smooth variation, the discontinuity could affect SpaNorm’s ability to separate real biology from the underlying unwanted variation. We are currently extending SpaNorm’s model to deal with and subsequently remove FOV effect. More generally, our approach for decomposing smooth spatial variation can be extended to accommodate other types of spatial omics data such as Imaging Mass Cytometry data [17], although it would likely require adaptation of the underlying models beyond the Negative Binomial distribution.

In conclusion, the development of both spot-based and subcellular spatial transcriptomics technologies is revolutionizing molecular biology. We identify strong library size variation across many ST datasets, which challenges standard normalization methods developed for scRNA-seq data. To address this, we introduced the first spatially-aware normalization approach that performs local regional library size adjustment, providing a level of flexibility that is a common limitation of many global adjustment approaches. We illustrate that our novel method outperforms the current state-of-the-art normalization methods, allowing a more accurate identification of spatially variable genes as well as regional detection. Furthermore, SpaNorm works equally well with segmented cell-level data and spot-based data, where each spot contains multiple cells. SpaNorm is available as an R package from https://github.com/bhuvad/SpaNorm

## Material and Methods

### SpaNorm model

To develop SpaNorm, a normalisation method that utilise spatial information while allowing optimal identification of spatial domains and spatially variable genes (SVGs), we model the count data using generalized linear model. Specifically, we assume that the count for gene *g* and spot (cell) *c* can be modelled as *z_gc_∼* Negative Binomial(NB)(*µ_gc_, ψ_g_*) where *ψ_g_*is the gene-specific dispersion parameter. The library size (LS) and biology affect the mean parameter through a log-linear model

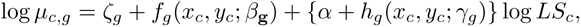

where (*x_c_, y_c_*) are the spatial coordinates and *LS_c_* is the library size for spot *c*. The two functions *f_g_*(*x_c_, y_c_*) and *h_g_*(*x_c_, y_c_*) are two-dimensional, gene-specific spatially-smooth functions constructed using 2D splines with *K* (default = 6) degree of freedom in each dimension, expressed as

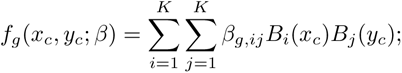

and

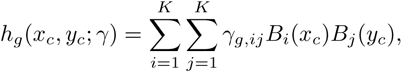

where *B_i_*(.) and *B_j_*(.) are B-splines basis functions and *α*+*h_g_*(*x_c_, y_c_*; *γ_g_*) can be thought as the gene- and location-specific scaling factor.

To improve fit, we also found that it is beneficial to ’regularise’ *β_g,ij_* and *γ_g,ij_* parameters using *L*_2_ penalty.

### Adjusted data

SpaNorm outputs a matrix of percentile-invariant adjusted count (PAC) that can be used for downstream analyses. For gene *g* and spot (cell) *c*, the PAC is calculated as quantile of a Negative Binomial distribution where the mean parameter does not contain library size effects,

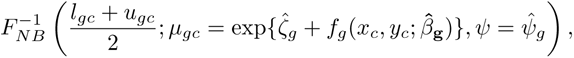

where

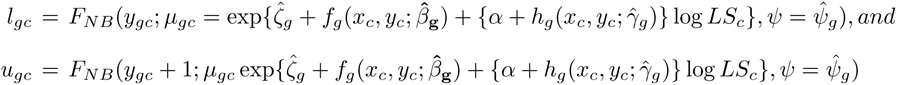

are the cumulative density functions of the Negative Binomial distribution which includes the library size effects. After obtaining the PAC, the log PAC was simply obtained as log(*PAC* + 1).

### Datasets

We use 6 datasets encompassing 27 samples (25 grid-based and 2 segmentation-based), four different platforms (Visium, Xenium, STOmics and CosMx), three tissues (Brain, Breast and Lung), and two species (Human and Mouse) to compare the performance of SpaNorm against no Normalisation and four other state-of-the-art normalisation approaches namely, Giotto, scran, RUV-III-NB and sctransform normalizations.

For the grid-based datasets, transcript detection tables for the 10x Xenium breast cancer dataset (IDC and ILC), 10x Xenium mouse brain, the NanoString CosMx non-small cell lung cancer, and the BGI STOmics mouse brain were obtained from [10]. Independently acquired region annotations were available from this dataset. These were obtained through image registration of DAPI images to reference tissue atlases, or through annotation of immunoflourescence or histology images. The 10x Visium human DLPFC dataset [15] was obtained through the SpatialLIBD R/Bioconductor package [18].

The two segmentation-based datasets (Xenium Human Breast Cancer Xenium datasets 1 and 2) were downloaded from https://www.10xgenomics.com/products/xenium-in-situ/preview-dataset-human-breast and subjected to further quality control (QC) steps that can be found in [13].

#### Data preprocessing

Measurements from all datasets, except the 10x Xenium breast cancer dataset with replicates, were allocated to regular hexagonal bins using the SubcellularSpatialData R/Bioconductor package. The *bins* parameter was set to 200 for the 10x Xenium breast cancer and mouse brain datasets, and 100 for the BGI STOmics and NanoString CosMx datasets. Bins where measurements spanned multiple regions were annotated based on the most frequent region annotation.

For the Xenium Human Breast Cancer Xenium datasets, segmentation was performed using BIDCell [13]. Default parameter values from the exemplar file for Xenium and the provided single-cell reference file were used (both files were downloaded from the official BIDCell repository). The model was trained end-to-end from scratch for 4000 iterations (i.e., using 4000 training patches). This amounted to a maximum of 22% of the entire image, thereby leaving the rest of the image unseen by the model during inference. Weights of the convolutional layers were initialised using He and colleagues’ approach [19]. We employed standard on-the-fly image data augmentation by randomly applying a flip (horizontal or vertical), rotation (of 90, 180, or 270 degrees) in the (*x*,*y*) plane. The order of training samples was randomised prior to training. We employed the Adam optimiser [20] to minimise the sum of all losses at a fixed learning rate of 0.00001, with a first moment estimate of 0.9, second moment estimate of 0.999, and weight decay of 0.0001.

#### Normalisation methods

Each dataset was normalised using the following methods:

- No Normalisation:Raw counts were log transformed. A pseudo count of 1 was added to all observations to avoid taking a logarithm of zero count.
- scran normalisation: A minimum size factor of 10*^−^*^8^ was imposed to avoid negative and zero size factor estimates. [6]
- sctransform normalisation [7].
- RUV-III-NB normalisation [11] with *K* = 1. Details of negative control features used and selection of pseudo-replicates can be found in [10].
- Giotto normalisation [8].
- SpaNorm normalisation (see above for details).

All normalisation methods were applied using their default parameters.

### Evaluation methods

#### Evaluating region-specific library size effects: annotation-based

For each dataset, we picked the top 1000 most abundant genes and fitted the following two negative binomial (NB) regression models for each gene:

- Model 1 (M1): Count *∼* AnnotatedRegion + log LS; and
- Model 2 (M2): Count *∼* AnnotatedRegion + log LS + log LS x smooth spatial term.

The Likelihood ratio test (LRT) was used to compare M1 vs M2 for each gene and the associated p-value was recorded. Then the proportion of genes for which M2 provides a significantly better fit was estimated using the qvalue function from qvalue Bioconductor package [21].

#### Evaluating region-specific library size effects: grid-based

Each dataset was split into rectangular grids. The size of the grids is dataset-specific because we require a minimum of at least 300 spots (cells) inside each grid. For grid *i* inside dataset *j*, we fitted the following NB model to the observed counts from cell *k* and gene *g*,

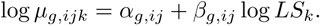

The heterogeneity of the library size effects (*β*’s) was tested for each gene using Cochran’s Q test [22].

#### Analysis of Variance

Oneway Analysis of Variance (ANOVA) was fitted to each gene with normalised data as a dependent variable and the manually-annotated regions as a factor (treatment) variable. The between-treatment and within-treatment variance estimates without and with a particular normalisation were compared in log-scale.

#### Simulation studies

We used scDesign3 pipeline [12] for simulating spatially variable genes (SVGs) (https://songdongyuan1994.github.io/scDesign3/docs/articles/scDesign3-DEanalysis-vignette.html) using Human DLPFC dataset 1 as the input dataset. To identify the true SVGs, two models were fitted: the first model contains both smooth spatial effects presenting the underlying biology and smoothly-varying library size effects, while the second model only contains the smoothly-varying library size effects. The deviance statistics of the two models were compared and genes with higher deviance differences were taken as containing a higher amount of smoothly-varying biology and the top 100 genes with the largest deviance difference were designated as the true SVGs.

#### Stably-expressed genes

We downloaded the list of stably-expressed genes for humans and mice from the Database of housekeeping genes and reference transcripts (https://housekeeping.unicamp.br/)[23].

#### Spatial domain identification

The spatial domain identification benchmark outlined in [10] was performed to study the impact of SpaNorm normalisation on spatial domain identification. Feature selection was performed on normalised datasets by identifying highly variable genes (HVGs). The top 1000, 2000, and 3000 genes were identified for datasets with genome-wide measurements. Where datasets were obtained using targeted panels, either genes with positive variance estimates from a fitted mean-variance trend, or all genes were selected. Dimensional reduction was performed using principal components analysis. Next, we used three clustering algorithms: the graph-based (Leiden, Louvain, or Walk-trap) algorithm from the igraph R package [24], BayesSpace [25], and SpaGCN [26] to perform clustering using the normalised data as input.

For graph-based algorithm, the graph was built using buildSNN function from the scran package [6] by setting the number of nearest neighbours to 10, 20, 30, or 50. For the Louvain and Leiden algorithms, 8 evenly spaced resolution parameters in the interval [0.1, 1] were assessed. BayesSpace and SpaGCN require the number of clusters to be pre-specified. As this is often unknown, we tested performance with the correct number of clusters, and over-/under-clustering by perturbing this number by 25%. SpaGCN was deployed from R using the reticulate and zellkonverter packages.

The defined parameter space was assessed exhaustively by running all possible combinations (27, 971, except a few failed runs). The CellBench framework was used to deploy the benchmark. The performance of the clustering algorithms to recover spatial domains under different normalisation strategies was compared by computing the Adjusted Rand Index (ARI) using the independently-annotated spatial regions as ground truth.

#### Spatially-variable genes (SVG) identification

MERINGUE [27] with default parameters was used to detect spatially variable genes. We chose MERINGUE over other methods because it takes normalised data as input. Hence it allows objective comparison of the impact of different normalisation on downstream analyses, in this case, SVG identification.

The strength of SVG signals was calculated using a statistic defined as 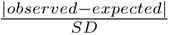. Finally, the concordance of these statistics between any two replicates of the same experiment was calculated using Spearman’s correlation coefficient.

#### Replicates used to calculate concordance

- Human DLPFC Visium set 1: Human DLPFC datasets 1-4
- Human DLPFC Visium set 2: Human DLPFC datasets 7-8
- Human DLPFC Visium set 3: Human DLPFC datasets 9-12
- Mouse Brain Xenium: Mouse Brain Xenium datasets 1-3
- Human NSCLC (Lung) CosMx: Human NSCLC (Lung) CosMx datasets 1-3
- Human BRCA Xenium: Human BRCA Xenium datasets 1-2

## Code availability

SpaNorm is implemented as an R package available from https://github.com/bhuvad/SpaNorm.

## Authors’ contributions

AS and DDB conceptualized the study with input from JYY and MJD. AS and DDB implemented the algorithm, developed the R package and performed the benchmarking studies. CC and PY performed the simulation studies and contributed to the benchmarking studies. AS wrote the first draft of the manuscript with input from JYY. All authors contributed to writing and approved the submitted version of the manuscript.

## Acknowledgements

The authors would like to thank Xiaohang Fu for processing the Xenium breast cancer datasets and Marni Torkel for assistance in the creation of Figure 1E.

## Notes

### Competing Interest Statement

The authors have declared no competing interest.

