## Supplementary Figures for "SpaNorm: spatially-aware normalisation for spatial transcriptomics data"

### SpaNorm: spatially-aware normalization for spatial transcriptomics data
### Supplementary Materials

Agus Salim *et al.*

May 31, 2024

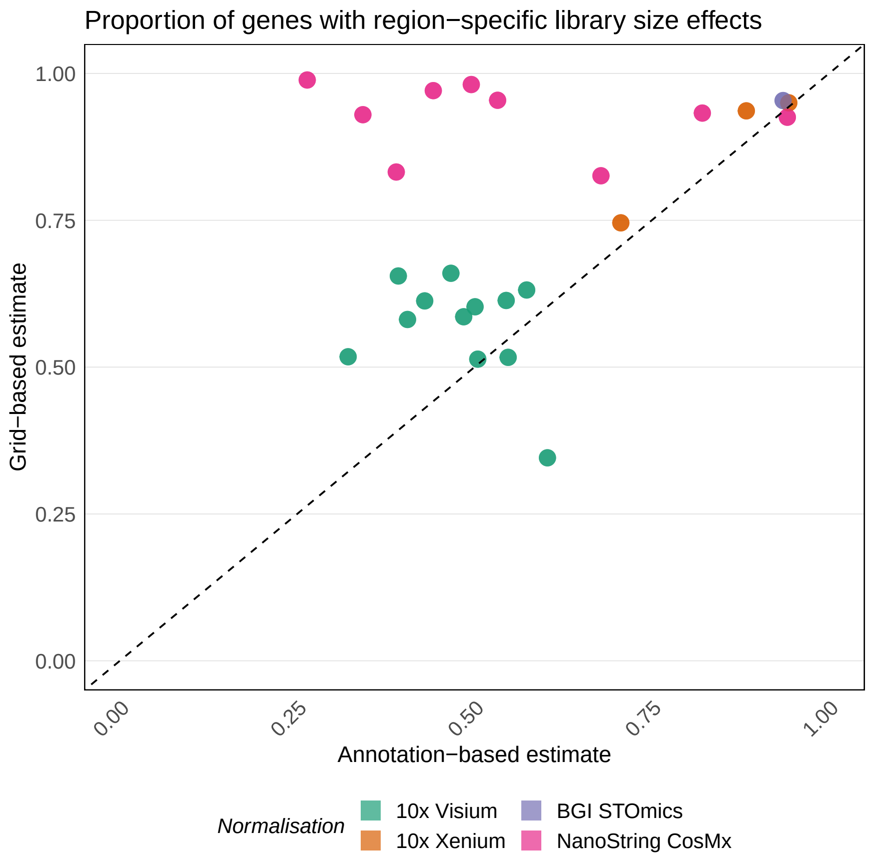


Supplementary Figure 1: Proportion of genes with region-specific library size effect using annotation-based model (x-axis) vs difference between annotation-based vs grid-based estimates (y-axis). Points above the diagonal line are datasets for which grid-based estimates are higher.

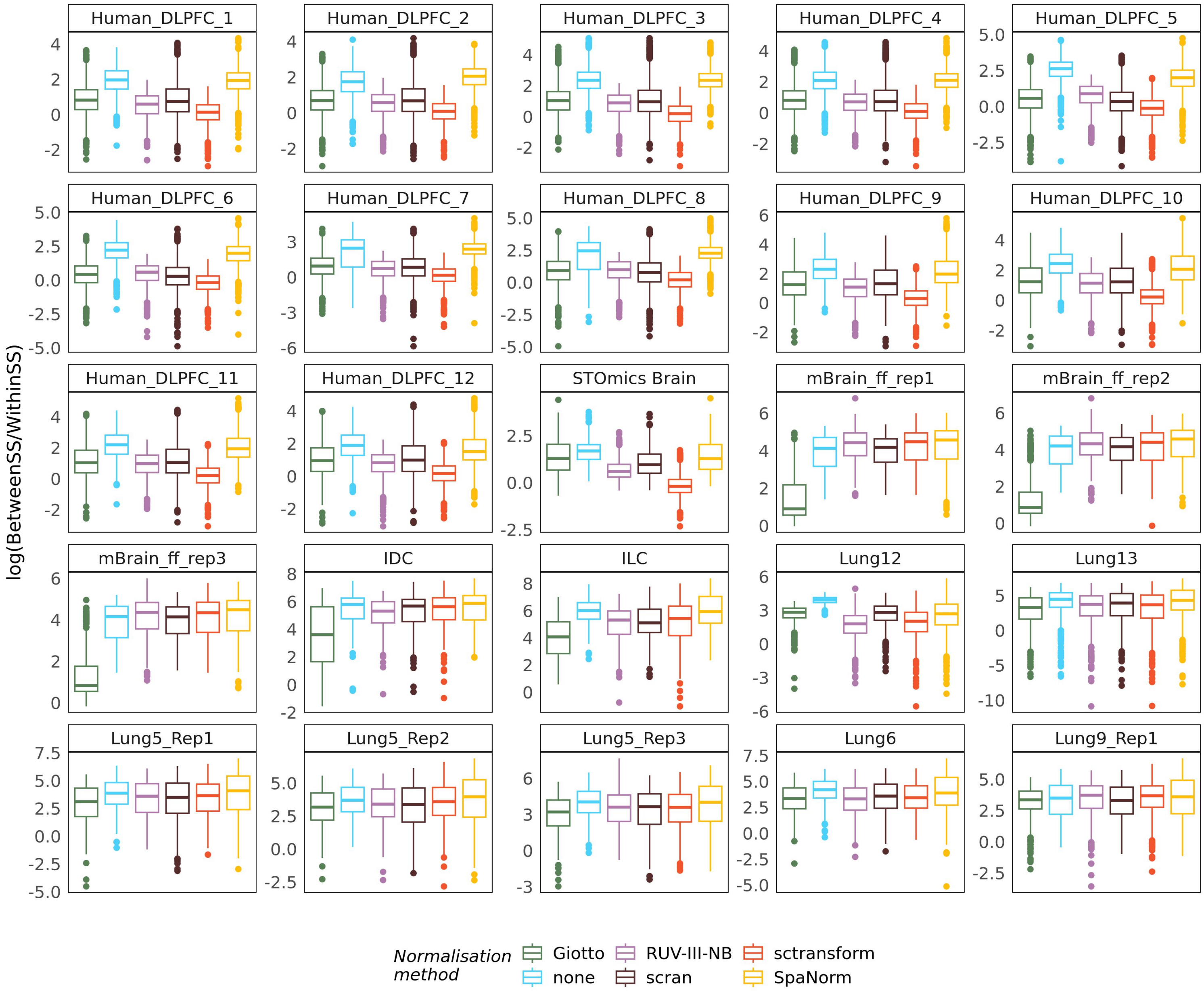


Supplementary Figure 2: Log ratio of (genewise) between-region to within-region variance in differently-normalised data. Higher log ratio indicates better between-region (spatial domain) signals. When compared to No-normalisation, SpaNorm is the only method that consistently improves spatial domain signals.

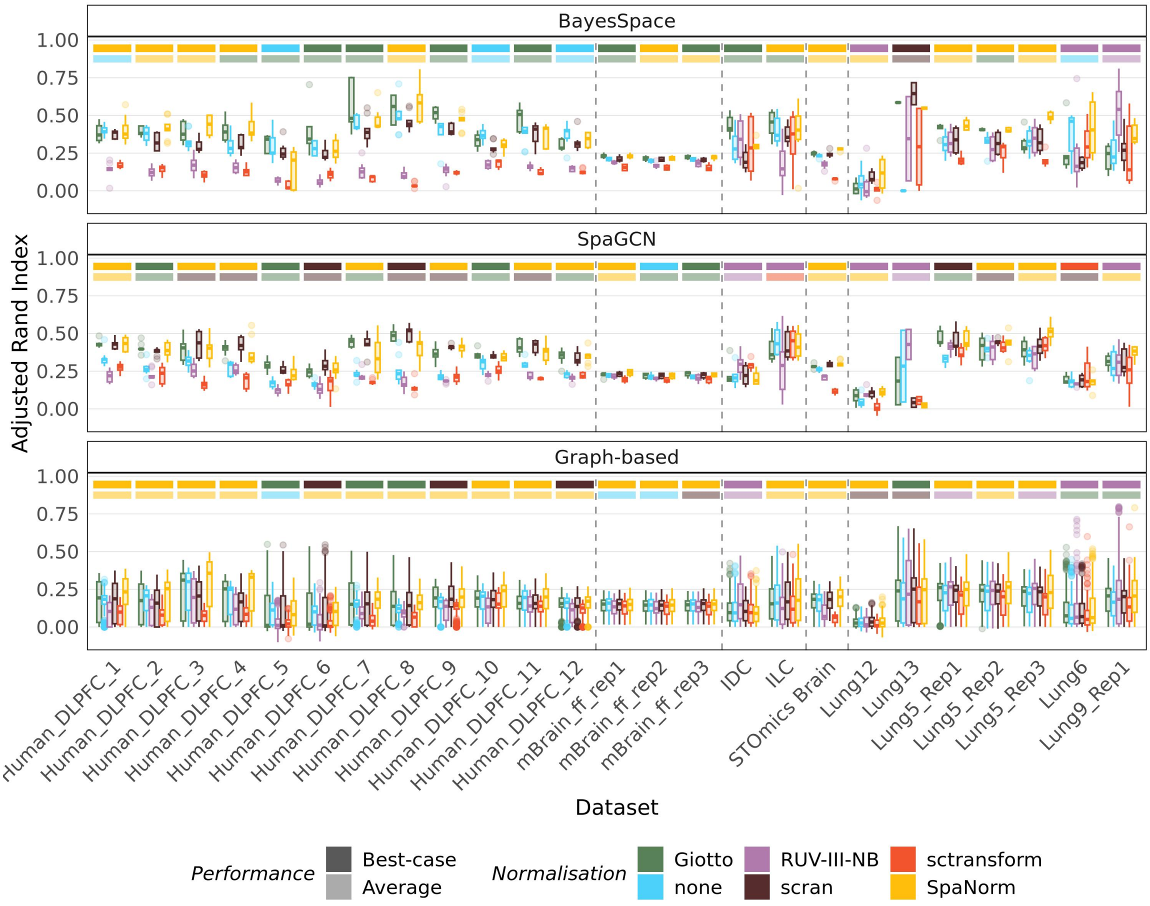


Supplementary Figure 3: Adjusted Rand Index of clusters identified using differently normalised data vs annotated spatial regions. The coloured bars above each group of boxplots indicate the best-performing methods based on maximum (darker-shade) and median (lighter-shade) statistic.

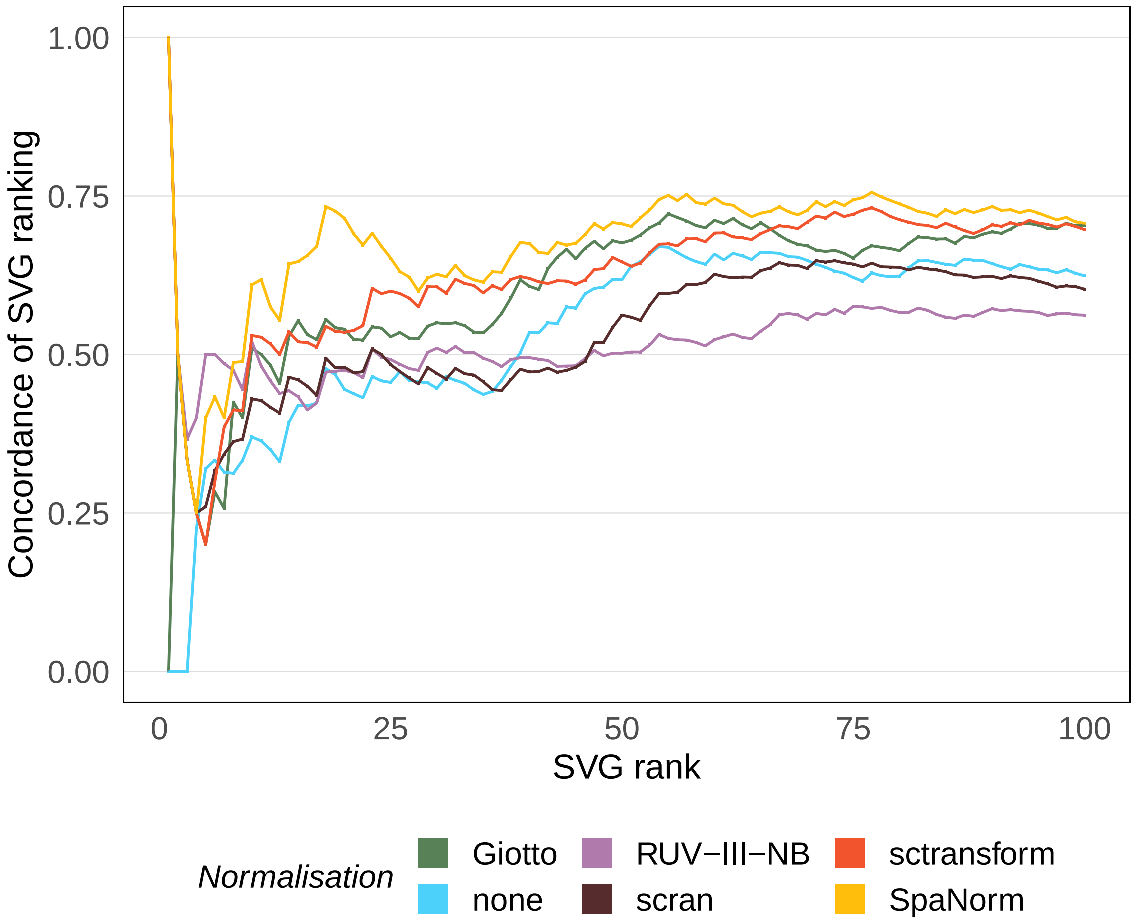


Supplementary Figure 4: Simulated datasets: The average proportion (from 10 simulated datasets) of the true spatially variable genes (SVGs) found among the top 100 SVGs identified by differently-normalised data. SpaNorm has the highest concordance followed by scran and Giotto.

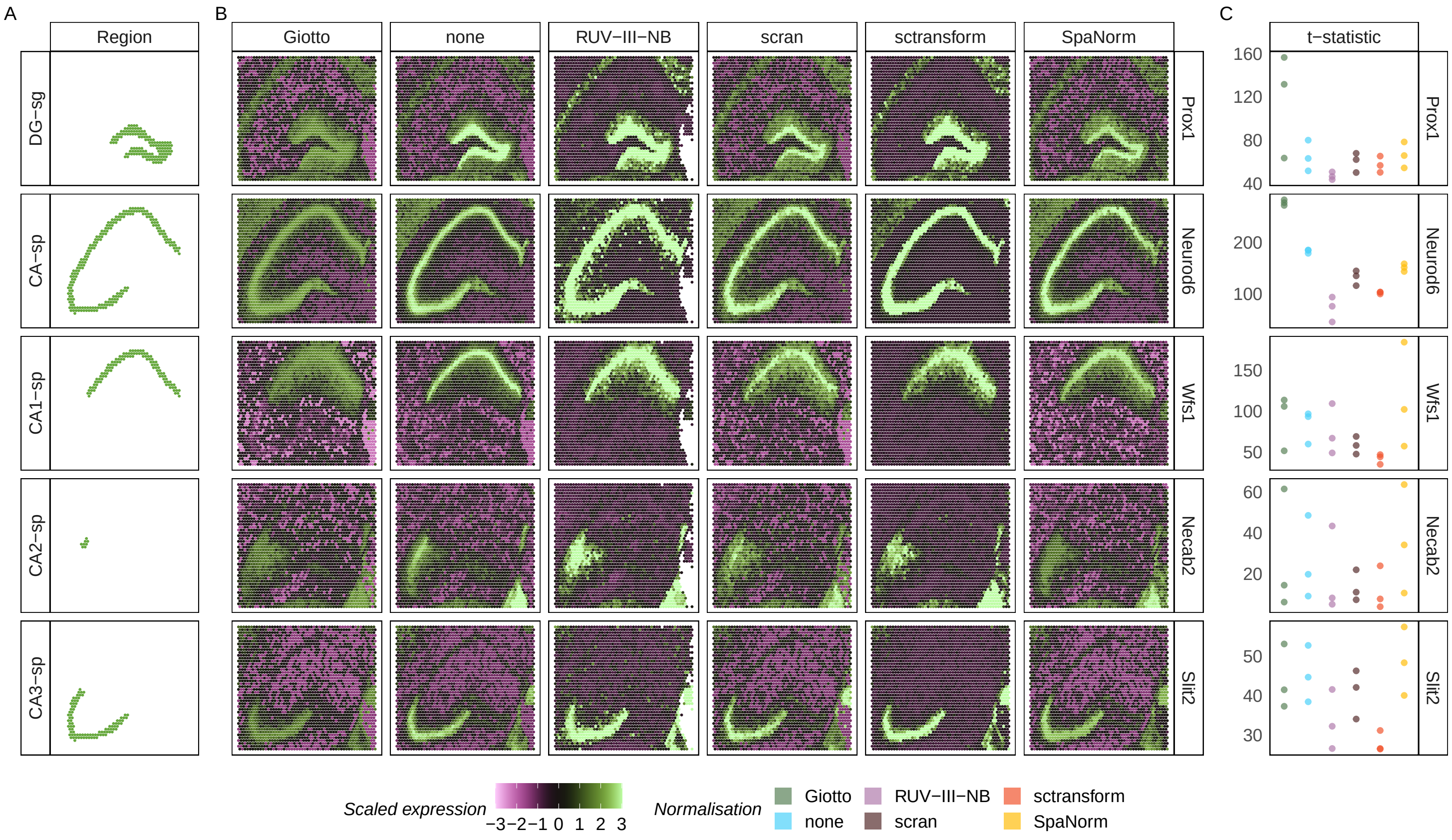


Supplementary Figure 5: Xenium mouse brain replicate 2. (A) Sub-structures of the mouse hippocampus. (B) Expression of 5 spatially variable genes (SVGs) in Xenium Mouse Brain dataset (replicate 1) under differently normalised data. *Prox1* differentiates the dentate gyrus (DG-sg) from pyramidal layers in CA1-3 (*Neurod6*). *Wfs1*, *Necab2 Slit2* are enriched in CA1-sp, CA2-sp and CA3-sp regions, respectively. (C) t-statistic from three replicates for comparing expression inside vs outside the regions. Higher statistic means stronger spatial signals.

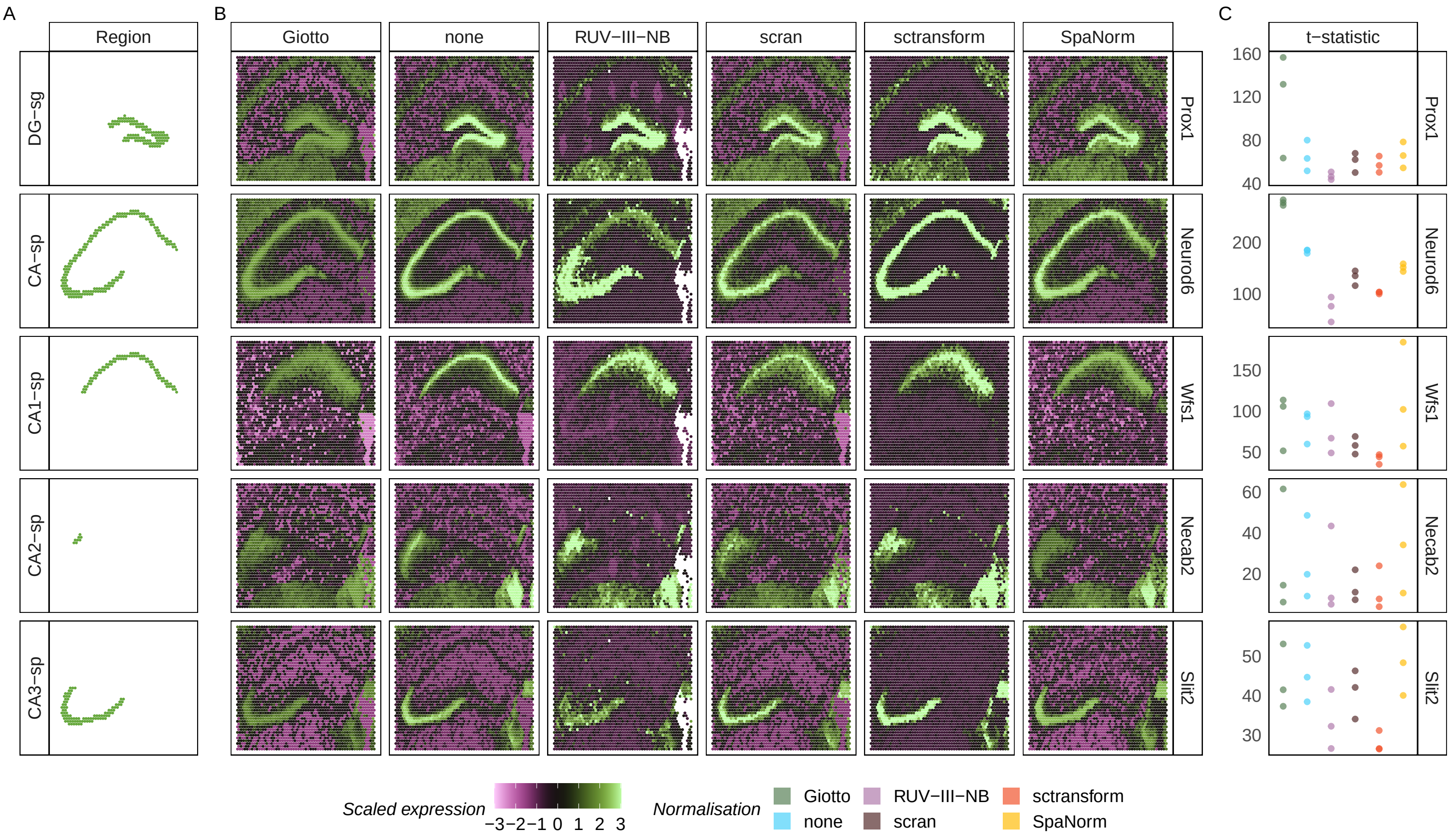


Supplementary Figure 6: Xenium mouse brain replicate 3. (A) Sub-structures of the mouse hippocampus. (B) Expression of 5 spatially variable genes (SVGs) in Xenium Mouse Brain dataset (replicate 1) under differently normalised data. *Prox1* differentiates the dentate gyrus (DG-sg) from pyramidal layers in CA1-3 (*Neurod6*). *Wfs1*, *Necab2 Slit2* are enriched in CA1-sp, CA2-sp and CA3-sp regions, respectively. (C) t-statistic from three replicates for comparing expression inside vs outside the regions. Higher statistic means stronger spatial signals.

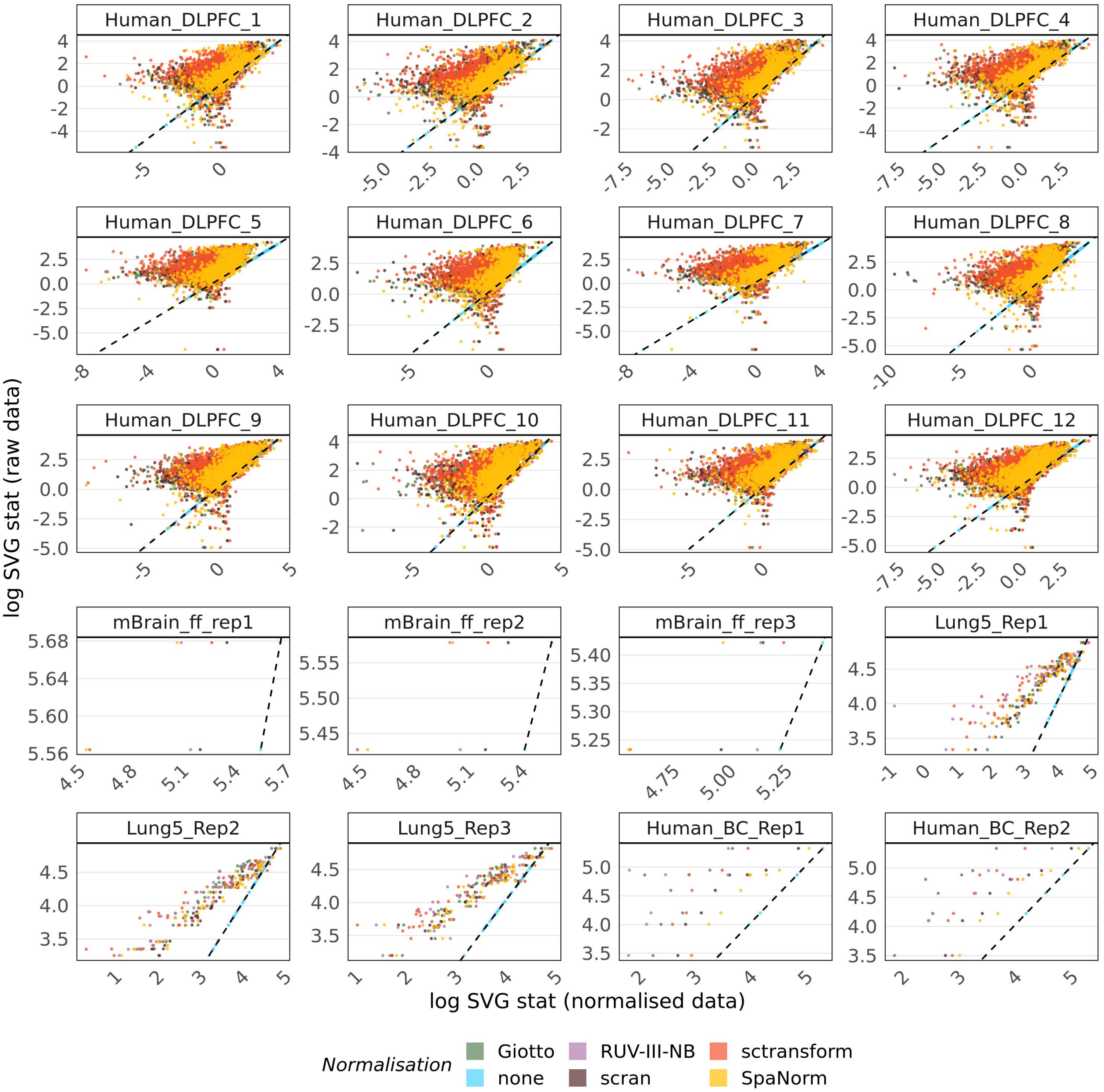


Supplementary Figure 7: Log of MERINGUE's SVG statistic with normalised data (x-axis) vs with raw data (No-normalisation) for genes that are stably expressed across cell types. Points above diagonal line indicates stronger SVG signals in the raw vs normalised data.

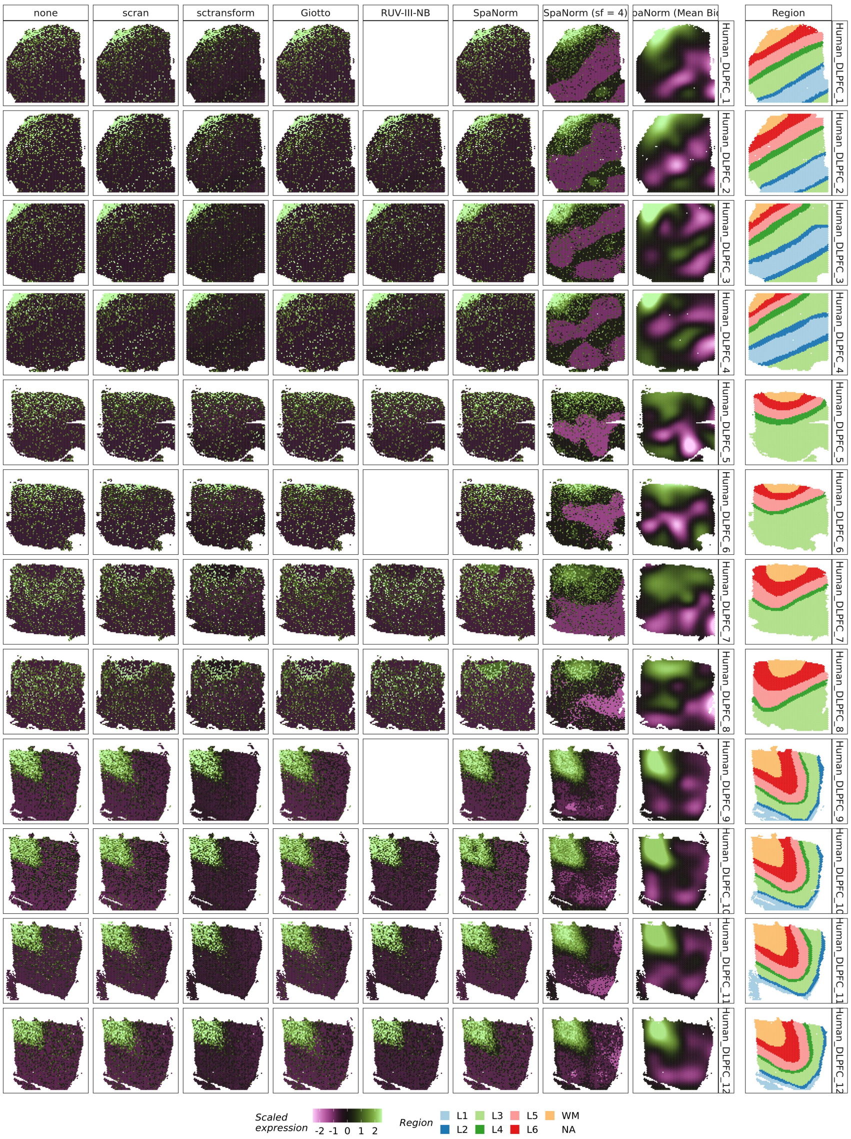


Supplementary Figure 8: Expression of MOBP, a marker of oligodendrocytes that are enriched in the white matter, following normalisation using different approaches. Expression of this gene is particularly low in samples 7 and 8, where most normalisation approaches are unable to recover signal. SpaNorm is able to identify this signal (Mean Bio), and can enhance the signal by scaling the counts (sf = 4).

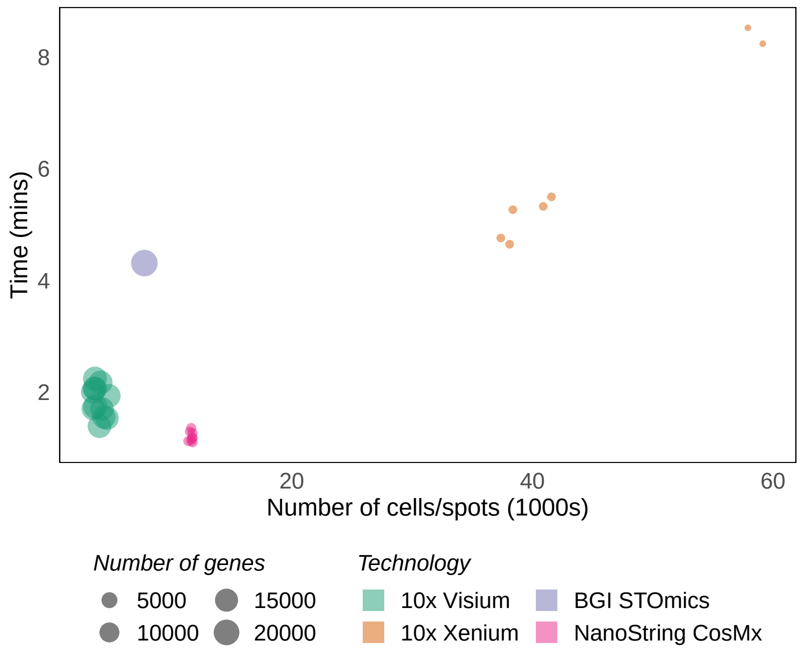


Supplementary Figure 9: Running time of SpaNorm on a Linux-based High Performance Computing (HPC) server with 1 core and 256Gb RAM
